# Structural basis for repurposing a 100-years-old drug suramin for treating COVID-19

**DOI:** 10.1101/2020.10.06.328336

**Authors:** Wanchao Yin, Xiaodong Luan, Zhihai Li, Leike Zhang, Ziwei Zhou, Minqi Gao, Xiaoxi Wang, Fulai Zhou, Jingjing Shi, Erli You, Mingliang Liu, Qingxia Wang, Qingxing Wang, Yi Jiang, Hualiang Jiang, Gengfu Xiao, Xuekui Yu, Shuyang Zhang, H. Eric Xu

**Affiliations:** The CAS Key Laboratory of Receptor Research, Shanghai Institute of Materia Medica, Chinese Academy of Sciences, Shanghai 201203, China; School of Medicine, Tsinghua University, Haidian District, Beijing, China; Department of Cardiology, Peking Union Medical College Hospital, Peking Union Medical College and Chinese Academy of Medical Sciences, Beijing, China; Tsinghua-Peking Center for Life Sciences, Tsinghua University, Beijing, China; Cryo-Electron Microscopy Research Center, Shanghai Institute of Materia Medica, Chinese Academy of Sciences, Shanghai 201203, China; State Key Laboratory of Virology, Wuhan Institute of Virology, Center for Biosafety Mega-Science, Chinese Academy of Sciences, Wuhan, Hubei, 430071, P. R. China; University of Chinese Academy of Sciences, Beijing 100049, China; WuxiBiortus Biosciences Co. Ltd., 6 Dongsheng West Road, Jiangyin 214437, China

**Author notes:** These authors contributed equally to this work (Wanchao Yin, Xiaodong Luan, Zhihai Li, Leike Zhang, Ziwei Zhou). Correspondence and requests for materials should be addressed to H. Eric Xu,; or Shuyang Zhang; or Xuekui Yu; or Gengfu Xiao.

## Abstract

The COVID-19 pandemic by non-stop infections of SARS-CoV-2 has continued to ravage many countries worldwide. Here we report the discovery of suramin, a 100-year-old drug, as a potent inhibitor of the SARS-CoV-2 RNA dependent RNA polymerase (RdRp) through blocking the binding of RNA to the enzyme. In biochemical assays, suramin and its derivatives are at least 20-fold more potent than remdesivir, the currently approved nucleotide drug for COVID-19. The 2.6 Å cryo-EM structure of the viral RdRp bound to suramin reveals two binding sites of suramin, with one site directly blocking the binding of the RNA template strand and the other site clash with the RNA primer strand near the RdRp catalytic active site, therefore inhibiting the viral RNA replication. Furthermore, suramin potently inhibits SARS-CoV-2 duplication in Vero E6 cells. These results provide a structural mechanism for the first non-nucleotide inhibitor of the SARS-CoV-2 RdRp and a rationale for repurposing suramin for treating COVID-19.

**One Sentence Summary:** Discovery and mechanism of suramin as potent SARS-CoV-2 RNA polymerase inhibitor and its repurposing for treating COVID-19.

## INTRODUCTION

Severe acute respiratory syndrome coronavirus 2 (SARS-CoV-2) has caused a global pandemic of coronavirus disease 2019 (COVID-19), with over 33.1 million infections and 998,000 death as reported on September 28 of 2020(Dong et al., 2020; Gorbalenya et al., 2020). SARS-CoV-2 is a positive-sense, single-stranded RNA virus. In addition to SARS-CoV-2, several related beta-coronaviruses, including SARS-CoV and Middle East respiratory syndrome coronavirus (MERS-CoV), are highly pathogenic, which infections can lead to severe acute respiratory syndrome exacerbated by loss of lung function and death. Compared to SARS-CoV and MERS-CoV, SARS-CoV-2 appears to have much higher capacity of human to human infections, which result in a rapidly growing pandemic(Sanche et al., 2020). Finding an effective treatment for SARS-COV-2 through drug repurposing is an urgent but unmet medical need.

Suramin (Figure 1A) is a century-old drug that has been used to treat African sleeping sickness and river blindness(Brun et al., 2010; Hawking, 1958). Suramin has also been shown to be effective in inhibiting the replication of a wide spectrum of viruses, including enteroviruses(Ren et al., 2014), Zika virus(Albulescu et al., 2017), Chikungunya(Albulescu et al., 2015) and Ebola viruses(Henß et al., 2016). The viral inhibition mechanism of suramin appears to be very diverse, including inhibition of viral attachment, entry, and release from host cells in part through interactions with viral capsid proteins(Wiedemar et al., 2020). Recently, suramin has been shown to inhibit SARS-CoV-2 infection in cell culture, which was proposed to act through preventing entry of the virus(Salgado-Benvindo et al., 2020). Here we report that suramin is a potent inhibitor of the SARS-CoV-2 RNA-dependent RNA polymerase (RdRp), an essential enzyme for the life cycle of virus. The potency of suramin is at least 20-fold more potent than remdesivir, the current FDA approved nucleotide drug for COVID-19. Cryo-EM structure reveals that suramin binds to the RdRp active site, blocking the binding of both RNA template and primer strands. These results provide a rationale for repurposing suramin for COVID-19 and a structural template to design next generation drugs of suramin derivatives.

**Figure 1.**
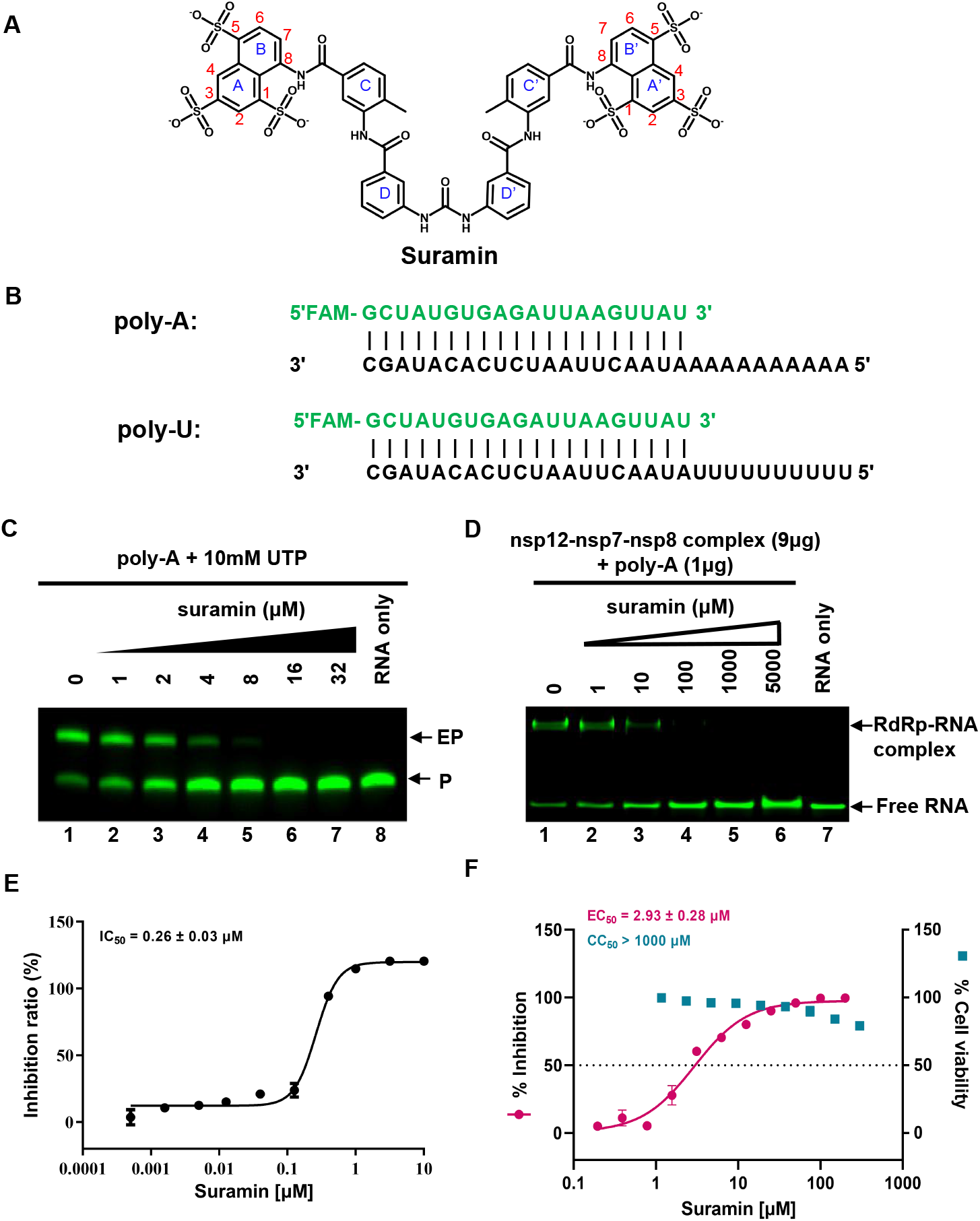
Inhibition of RdRp by suramin and the potential anti-SARS-CoV-2 effect of suramin. A. The chemical structure of suramin. B. The 30-base template and 20-base primer duplex RNAs with a FAM at the 5’ of the primer. Poly-A was used in the gel-based elongation assay, while Poly-U was used in the fluorescence- based activity assay for SARS-CoV-2 RdRp. C. Elongation of partial RNA duplex by the purified RdRp complex and its inhibition by suramin. The EP means the elongated product, while the P means the primer RNA strand. D. Gel mobility shift of the RdRp-RNA complex and the effect of suramin. E. IC_50_ of suramin for RdRp complex. F. EC_50_ of suramin for SARS-CoV-2 inhibition and CC_50_ of suramin for cell-based toxicity.

## RESULTS and DISCUSSION

### Inhibition of RdRp by suramin and the potential anti-SARS-CoV-2 effect of suramin

The core RNA polymerase of SARS-CoV-2 is composed of non-structural protein nsp12 with two accessary subunits nsp7 and nsp8(Kirchdoerfer and Ward, 2019; Subissi et al., 2014). Incubation of the purified nsp12-7-8 complex (Figure S1A-1B) with a 30-base template and 20-base primer (poly-A in Figure 1B) allowed the primer extension to the same length as the template in the presence of saturated concentrations of UTP as illustrated in a gel-based assay (lane 1 in Figure 1C). Addition of 8-32 μM suramin nearly abolished the elongation of the primer strand while it required 100-1000 μM of remdesivir in its triphosphate form (RDV-TP) to achieve the same degree of inhibition under the same condition(Yin et al., 2020). Interestingly, addition of 100 μM of higher concentrations of suramin completely blocked the formation of RdRp-RNA complex, while it required more than 5 mM of RDV-TP to inhibit the binding of RdRp to RNA (Figure 1D and Figure S1C). Solution based assays of the RdRp inhibition determined that half maximal inhibition concentration (IC_50_) of suramin is 0.26 μM (Figure 1E), and the IC_50_ for RDV-TP is 6.21 μM under identical assay conditions (Figure S1D), suggesting that suramin is at least 20-fold more potent than RDV-TP. Cell-based experiments indicated that suramin was able to inhibit SARS-CoV-2 duplication in Vero E6 cells with a half maximal effective concentration (EC_50_)of ~2.9 μM, which is about the same range as remdesivir in the same assay (Figure 1F and Fig. S1E)(Wang et al., 2020). The apparent weaker inhibition of suramin in cell-based assays than in enzyme inhibition assays may be due to the highly negative charges of suramin that may prevent its efficient uptake by the host cells. In addition, the CC_50_ (concentrations of drug required to reduce cell viability by 50%) of suramin is over 1000 μM, indicating its relatively low cytotoxicity, which is much safer than remdesivir (Figure 1F and Fig. S1E).

### The structure of the RdRp-suramin complex

For the cryo-EM studies, we incubated the SARS-CoV-2 RdRp complex with 10-fold molar excess of suramin (see methods). The structure was determined at a nominal resolution of 2.57 Å with 95,845 particles from over 8 million original particles auto-picked from 11,846 micrographs (Figure S2 and Table S1). Because of the relatively high resolution of the structure, the EM map reveal the clear density for all key components of the RdRp-suramin complex, including one nsp12 (residues S6-C22, V31-I106, M110-L895, and N911-T929), one nsp7 (residues K2-G64), two nsp8 (residues D78-A191 for nsp8-1, and residues T84-A191 for nsp8-2, respectively), and two suramin molecules (Figure 2A and Figure S3).

**Figure 2.**
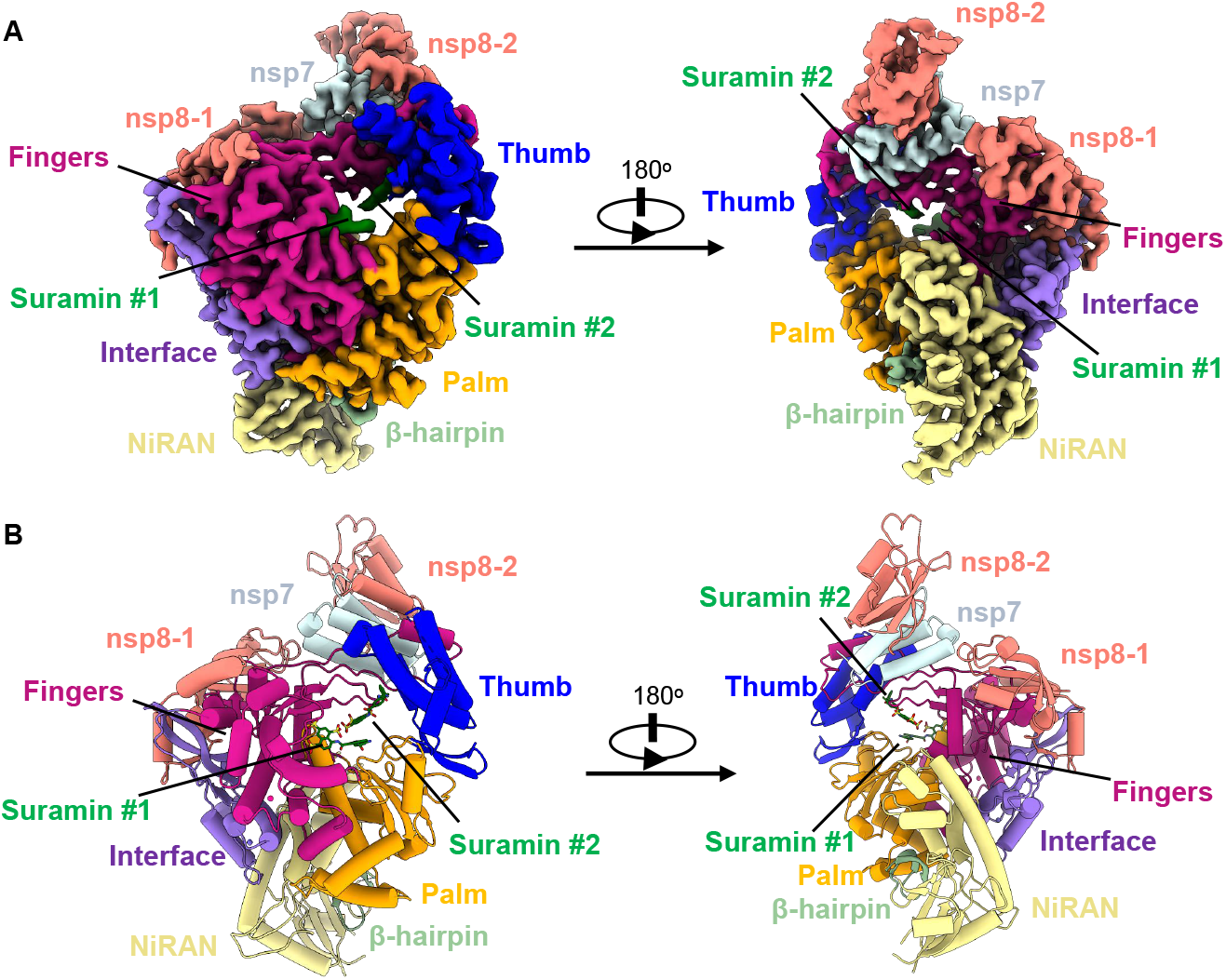
Overall Structures of the RdRp-suramin complex. A. Two views of cryo-EM density map of the RdRp-suramin complex, SARS-CoV2 nsp12 contains a N-terminal extension composed of the β-hairpin (green), the NiRAN domain (yellow) and an interface domain (purple) adjacent to the RdRp core domain, which contains subdomains: fingers, palm, and thumb, colored with violet, orange, and blue, respectively. The nsp12 binds to a heterodimer of nsp7 (light green) and nsp8 (nsp8-2, red) as well as to a second subunit of nsp8 (nsp-1, red). The two bound suramin molecules are set as dark green. Color code is used throughout. B. Two views of the overall structure of the RdRp-suramin complex, the color scheme is according as above.

The overall structure of the RdRp-suramin complex is very similar to the apo RdRp complex, with an RMSD of 0.465 Å for all Cα atoms between the two structures (Figure 2B and Figure S4). Nsp12 adopts the same right-hand palm-fingers configuration, where its catalytic active site is composed of seven highly conserved motifs A-G (Figure 3A). Two suramin molecules were found to fit into the catalytic chamber (Figure 2A-2B, and Figure 3B).

**Figure 3.**
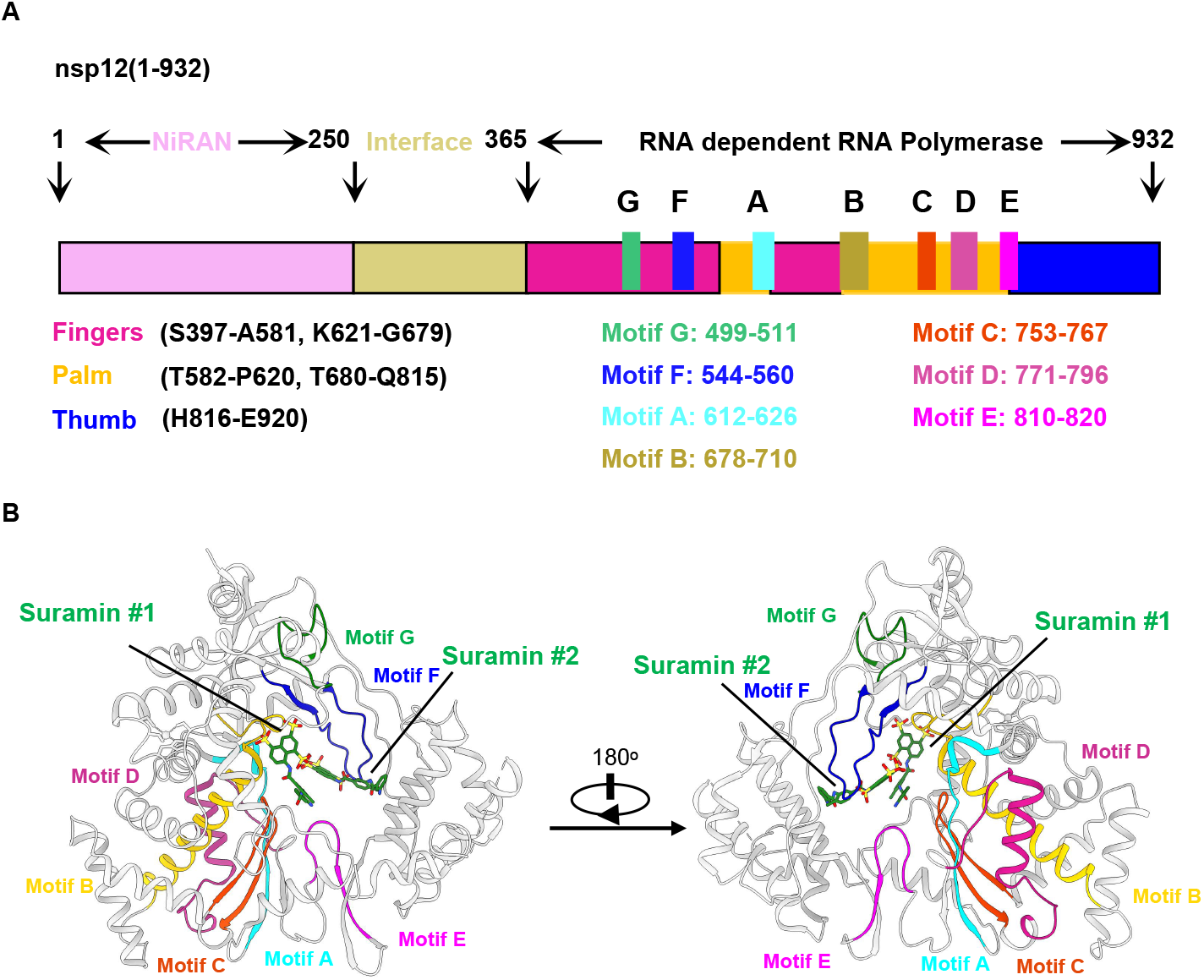
The conserved motif A-G in the RdRp-suramin complex. A. Schematic diagram for the components of the RdRp complex subunit nsp12, the motif (A to G) are highlighted. B. The color code is as follows: motif A (cyan), motif B (gold), motif C (orange red), motif D (medium violet red), motif E (magenta), motif F (blue), and motif G (green).

### The interactions of suramin with SARS-CoV-2 RdRp

One suramin molecule (suramin #1) is fit into a cavity formed by conserved motif G and the N-terminus of motif B (Figure 4A and Figure 3). The chemical structure of suramin has a two-fold symmetry with a urea linker at the center (Figure 1A). The EM density map for this suramin is very clear defined but only for half of the suramin molecule without the urea linker (Figure 4A and 4B). The key interactions of suramin #1 with RdRp were summarized in Figure 4C and Table 1, including hydrogen bonds, charge interactions, and hydrophobic packing interactions with conserved RdRp residues, which restrain the naphthalene-trisulfonic acid head in a relative narrow cavity. Two out of the three sulfonates (positions 3 and 5) form hydrogen bonds with the side chains from N497, K500, R569 and Q573, and the main chain from N497, while the sulfonate at positions 1 points toward the solvent and forms only one hydrogen bond with the side chain of N496. The K577 side chain forms cation-π stacking with the naphthalene ring, and also forms a hydrogen bond with the amide bond linker between the naphthalene and benzene rings. The amide bond linker between the benzene rings C and D forms a hydrogen bond with main chain NH of G590. In addition, the suramin # 1 is in van der Waals contact with several residues, including L576, A580, A685, Y689 and l758. The other suramin molecule (suramin #2) is fit into the cavity near the catalytic active site formed by conserved motifs A, C, E, and F (Figure 4B and Figure 3). Also, only half of suramin was observed in the structure with clear EM density map. The key interactions of suramin #2 with RdRp were summarized in Figure 4D and Table 1, including hydrogen bonds, charge interactions, and hydrophobic packing interactions. Different from suramin # 1, the sulfonate at positions 5 of suramin # 2 points toward the solvent and forms only one hydrogen bond with the side chain of R555, while the other two sulfonates at positions 1 and 3 form hydrogen bonds with the side chains from K551, R553, R555 and R836, and the main chains from A550 and K551. Meanwhile the side chain of R555 also forms a hydrogen bond with the amide bond linker between the naphthalene and benzene rings. The R836 side chain forms cation-π stacking with the benzene ring C. The NH of the benzene ring D forms a hydrogen bond with the side chain of D865. In addition, suramin # 2 is in van der Waals contact with several residues, including H439, I548, S549, A840, S861 and L862. Sequence alignment with RdRp from several viruses indicated that these suramin-contacting residues are conserved (Figure S5), suggesting that suramin may be a general inhibitor of viral RdRp, which implies that suramin could be used as a drug for treating infections of these viruses.

**Table 1.**
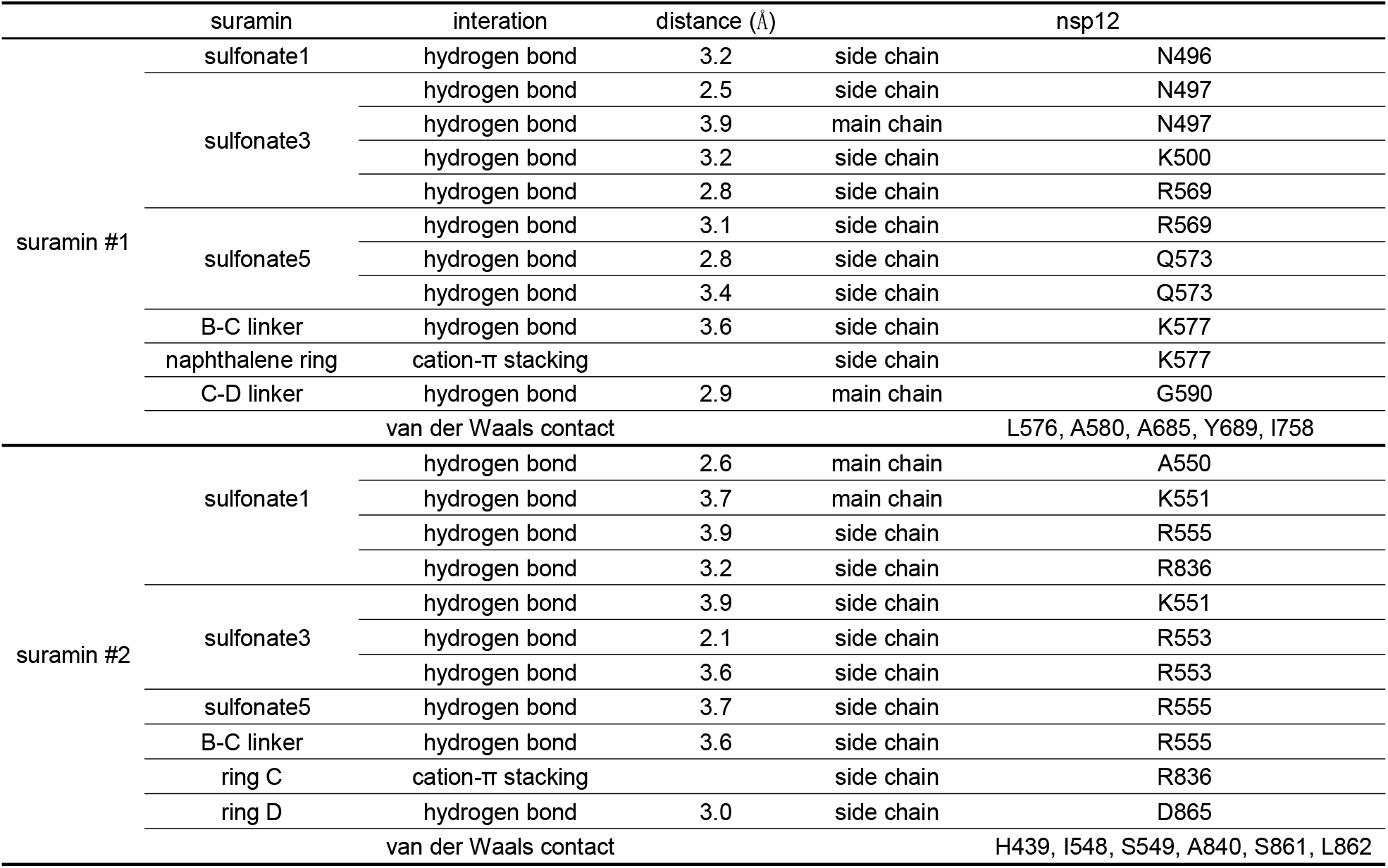
Interactions between suramin and nsp12 in the RdRp-suramin structure.

**Figure 4.**
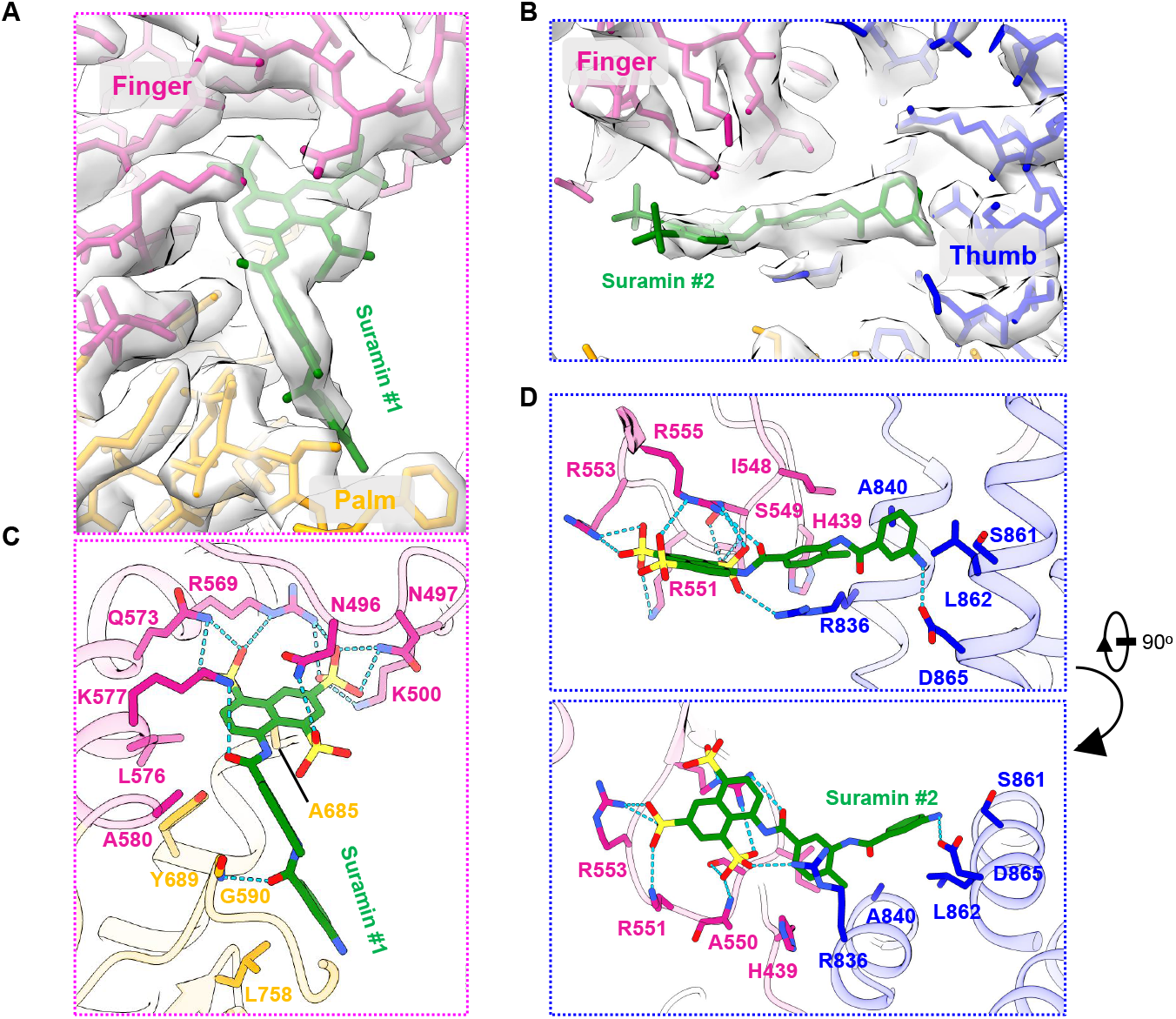
Close views of the interactions between the SARS-CoV-2 RdRp and suramin. A-B. The maps for the two suramin molecules. C-D. Interactions of the two suramin molecules with RdRp. The hydrogen bond is displayed as green dashed line.

### Inhibition mechanism of suramin to the SARS-CoV-2 RdRp

Structural comparison of the RdRp-suramin complex with the remdesivir-bound RdRp complex reveal a clear mechanism of RdRp inhibition by suramin (Figure 5A). If the base position of remdesivir was defined as +1 position, then the binding of the first suramin molecule occupies the same space of −1 to −3 positions of RNA template strand (suramin #1 in Figure 5B). The second suramin molecule at the active site occupies the space of the primer strand ranging from −4 to +1 positions (suramin #2 in Figure 5C). The binding of these two suramin molecules would thus block the binding of the RNA template-primer duplex to the active site as well as the entry of nucleotide triphosphate into the catalytic site, which results in the direct inhibition of the RdRp catalytic activity. The direct inhibition mechanism of SARS-CoV-2 RdRp by suramin is different from the suramin-mediated inhibition of the norovirus RdRp, which also contained two binding sites for suramin binding(Mastrangelo et al., 2012). In each site, only half of suramin molecule was found. Structural comparison of the SARS-CoV-2 RdRp with the norovirus RdRp reveals only one of the two suramin binding sites (suramin #2) is partially overlapped (Figure S6A, S6C and S6D). Both suramin binding sites in norovirus RdRp did not clash with the RNA strands but one of the suramin binding site is overlapping with the proposed nucleotide entry channel, thus indirectly blocking RdRp polymerization activity, a mechanism different from the direct blocking of the binding of the RNA template to the SARS-CoV-2 RdRp by suramin (Figure 5). In addition, structural comparisons of the SARS-CoV-2 RdRp-suramin structure with the structures of the norovirus RdRp bound to other suramin derivatives show that the suramin and suramin derivatives bind to the RdRp with diverse conformations and orientations(Croci et al., 2014; Mastrangelo et al., 2012) (Figure S6B, S6E and S7), indicating the possibility for other suramin derivatives to inhibit the SARS-CoV-2 RdRp as tested below.

**Figure 5.**
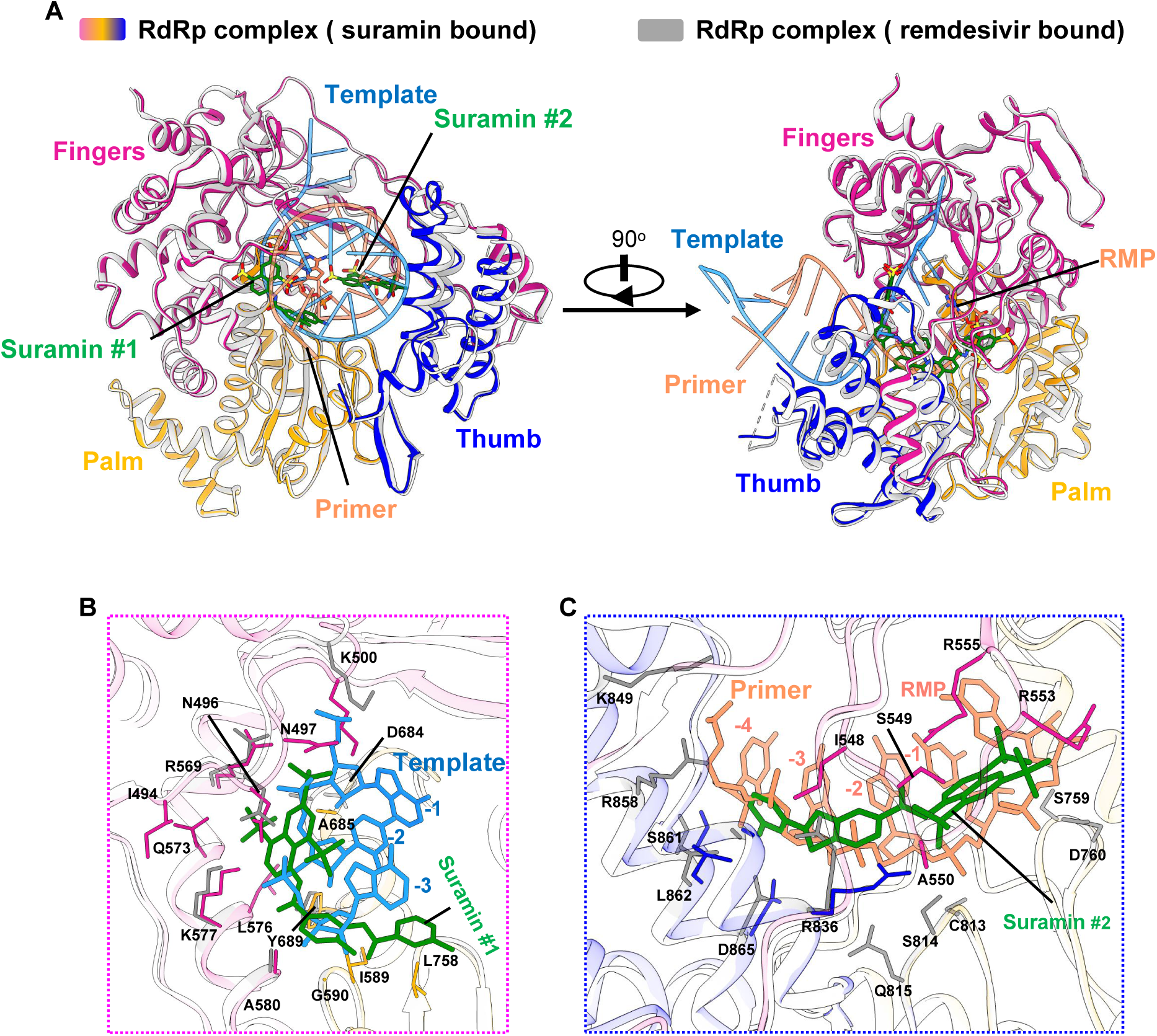
Inhibition mechanism from comparison with the remdesivir-bound RdRp structure. A. The two overall views of the RdRp-suramin complex overlapped with the remdesivir-bound RdRp structure (PDB ID: 7BV2). For clarity, only the polymerase domains are presented. The remdesivir bound RdRp structure is set as light gray, the template RNA is set as cyan, and the primer RNA is set as red. B. Close view of suramin #1 overlapped with RNA template strand. C. Close view of suramin #2 overlapped with RNA primer strand.

### Inhibition SARS-CoV-2 RdRp by suramin derivatives

Suramin derivatives have been explored for diverse applications including anti-cancers and anti-parasites(Wiedemar et al., 2020). To determine the structure-activity-relationship (SAR), we screened a set of 8 different suramin derivatives using in vitro RdRp primer extension assays (Figure 6A and Figure S1F). All 8 tested suramin derivatives showed efficient inhibition of RdRp activity (Figure S1G). NF157, NF279, and NF449 are most potent inhibitors with IC_50_ of 0.05 μM, which are about 5-fold more potent than the parent drug suramin (Figure 6B). Cell-based assays showed that NF110 inhibited SARS-CoV-2 replication with EC_50_ of 2.87 μM (Figure 6C), while NF157 and NF279 inhibited SARS-CoV-2 replication with EC_50_ of ~10 μM. The CC_50_ values of all suramin derivatives are over 1000 μM, indicating their good safety window and their potential use for treating COVID-19. However, there is a 200-fold separation between their biochemical potency in inhibiting RdRp activity and their potency in inhibiting viral replication in cell-based assays, suggesting the difficulties of these suramin derivatives to be uptake by host cells(Alsford et al., 2013). Future drug formulation such as with glycol chitosan-based nanoparticles(Cheng et al., 2019) may improve their bioavailability to lung tissues and their potency in inhibiting viral replication.

**Figure 6.**
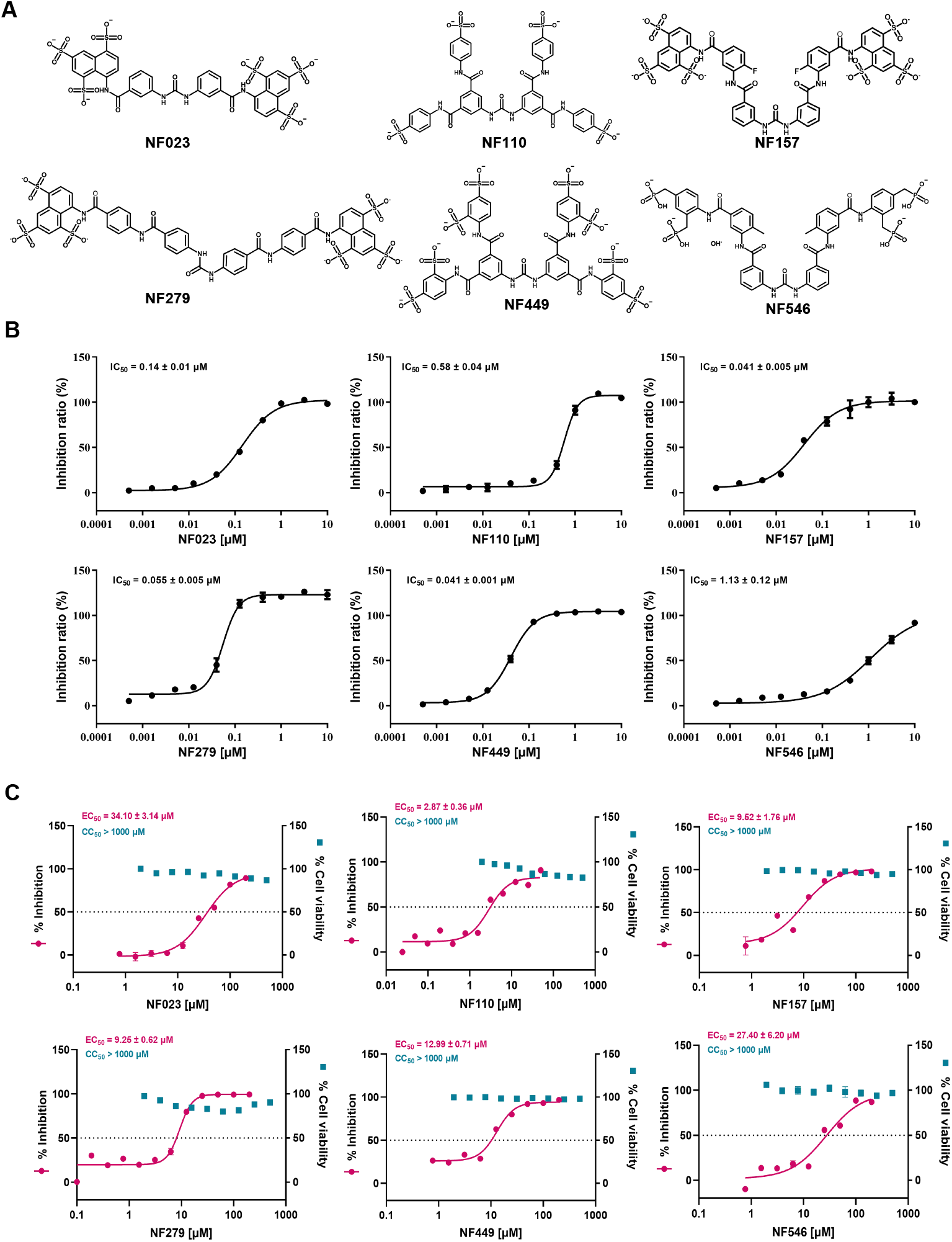
Inhibition SARS-CoV-2 RdRp by suramin derivatives. A. The chemical structures of suramin derivatives. B. The inhibition of RdRp activity by suramin derivatives. C. Inhibition of viral replication and cellular toxicity of suramin derivatives.

The COVID-19 pandemic has infected many more people and caused many more deaths than anyone could imagined since the outbreak of SARS-CoV-2 in December of 2019. The ongoing pandemic urgently press the need for effective vaccines and drug treatments. Suramin is a century-old drug that has been used for treating African sleepiness and showed activity against a number of viruses without a clear mechanism. In this paper, we showed that suramin is a direct potent inhibitor of the viral RdRp, the essential enzyme for the viral life cycle. The structure reveals that suramin binds to the active site of RdRp, blocking the binding of both strands of the template-primer RNA substrate and inhibiting the polymerase activity of the RdRp. Suramin derivatives also showed potent inhibition of the RdRp activity and blocked viral replication in cell-based assays. Together, these results uncovered the structural mechanism of the first non-nucleotide inhibitor of the SARS-CoV-2 RdRp and suggested the potential use of suramin for treating the SARS-CoV-2 infections. The structure and biochemical results presented in this paper also provided a rationale to develop suramin analogs as well as drug formulations to improve their potency and efficacy to inhibit SARS-CoV-2 replication.

## Supporting information

map_and_model

## Acknowledgments

The cryo-EM data were collected at the Cryo-Electron Microscopy Research Center, Shanghai Institute of Materia Medica. This work was partially supported by National Key R&D Program of China (2020YFC0861000), CAMS Innovation Fund for Medical Sciences No. 2020-I2M-CoV19-001, and Tsinghua University-Peking University Center for Life Sciences 045-160321001 to S. Z.; the National Key R&D Programs of China 2018YFA0507002; Shanghai Municipal Science and Technology Major Project 2019SHZDZX02 and XDB37030103 to H.E.X.; the 100 Talents Program of the Chinese Academy of Sciences, Chinese Academy of Sciences grant (XDA12010317), Natural Science Foundation of Shanghai (18ZR1447700) to X.Y; the National Natural Science Foundation of China (31900869) and Shanghai Sailing Program (19YF1456800) to Z.L; Science and Technology Commission of Shanghai Municipal 20431900100 and Jack Ma Foundation 2020-CMKYGG-05 to H.J.; National Natural Science Foundation 31770796, National Science and Technology Major Project2018ZX09711002, and K.C. Wong Education Foundation to Y.J, the National Natural Science Foundation of China (31970165) to L. Z.; the National Natural Science Foundation of China (81903433) to J. S..

## Author contributions

W.Y. designed the expression constructs, purified the RdRp complex, prepared samples and cryo-EM grids, performed data collection and processing toward the structure determination, analyzed the structures, and prepared figures and manuscript. X.L. designed RdRp activity assays and discovered RdRp inhibition by suramin, participated in data interpretation and figure preparation; Z.L., Z.Z., Q. W., and X.Y. evaluated the specimen by negative-stain EM, screened the cryo-EM conditions, prepared the cryo-EM grids and collected cryo-EM images, performed data processing, density map calculations, model building, and figure preparation; L.Z., Q.W., and G.X. designed and performed cell-based viral inhibition assays; X. W., F. Z., and M.G. participated in expression, purification and functional assays of the RdRp; J. S., E. Y. and M. L. participated in the analysis of suramin derivatives; Y.J. participated in experimental design and manuscript editing; H.J. conceived and coordinated the project; S.Z. conceived the project, initiated collaboration with H.E.X., and supervised X.L.; H.E.X. conceived and supervised the project, analyzed the structures, and wrote the manuscript with inputs from all authors.

## Data and materials availability

Density maps and structure coordinates have been deposited with immediate release. The accession numbers of Electron Microscopy Database and the Protein Data Bank are EMD-30572and PDB ID 7D4F for the SARS-CoV-2 RdRp bound to suramin. These files are immediately available for download as supplemental materials in this article. Materials are available up on request.

## Supplementary Material

### Materials and Methods

#### Constructs and expression of the RdRp complex

The RdRp complex was prepared according to same method reported (Kirchdoerfer and Ward, 2019) as described below. The full-length gene of the SARS-CoV-2 nsp12(residues 1-932) was chemically synthesized with codon optimization (General Biosystems). The gene was cloned into a modified pFastBac baculovirus expression vector containing a 5’ ATG starting sequence and C-terminal Tobacco Etch Virus (TEV) protease site followed by a His8 tag. The plasmid contains an additional methionine at the N-terminus and GGSENLYFQGHHHHHHHH at the C-terminus of nsp12. The full-length genes for nsp7 (residues 1-83) and nsp8 (residues 1-198) were cloned into the pFastBac vector containing a 5’ ATG starting sequence. All constructs were generated using the Phanta Max Super-Fidelity DNA Polymerase (Vazyme Biotech Co.,Ltd) and verified by DNA sequencing. All constructs were expressed in *Spodoptera frugiperda* (S*f*9) cells. Cell cultures were grown in ESF 921 serum-free medium (Expression Systems) to a density of 2-3 million cells per ml and then infected with three separate baculoviruses at a ratio of 1:2:2 for nsp12, nsp7 and nsp8 at a multiplicity of infection (m.o.i.) of about 5. The cells were collected 48 h after infection at 27 °C and cell pellets were stored at – 80 °C until use.

In addition, the genes of nsp7 and nsp8 were cloned into a modified pET-32a (+) vector containing a 5’ ATG starting sequence and C-terminal His8 tag with a TEV cleavage site for expression in E. coli. Plasmids were transformed into BL21(DE3) (Invitrogen). Bacterial cultures were grown to an OD600 of 0.6 at 37 °C, and then the expression was induced with a final concentration of 0.1 mM of isopropyl β-D-1-thiogalactopyranoside (IPTG) and the growth temperature was reduced to 16 °C for 18-20 h. The bacterial cultures were pelleted and stored at – 80 °C until use.

#### Purification of the RdRp complex

The purification of nsp7 and nsp8 expressed in bacteria was similar to the purification of nsp7 and nsp8 reported previously(Kirchdoerfer and Ward, 2019). Briefly, bacterial cells were lysed with a high-pressure homogenizer operating at 800 bar. Lysates were cleared by centrifugation at 25,000 × g for 30 min and were then bound to Ni-NTA beads (GE Healthcare). After wash with buffer containing 50 mM imidazole, the protein was eluted with buffer containing 300 mM imidazole. The tag was removed with incubation of TEV protease overnight and protein samples were concentrated with 3 kDa or 30 kDa molecular weight cut-off centrifuge filter units (Millipore Corporation) and then size-separated by a Superdex 75 Increase10/300 GL column in 25 mM HEPES pH 7.4, 200 mM sodium chloride, 5% (v/v) glycerol. The fractions for the nsp7 or nsp8 were collected, concentrated to about 10 mg/ml, and stored at – 80 °C until use.

The insect cells containing the co-expressed RdRp complex were resuspended in binding buffer of 25 mM HEPES pH 7.4, 300 mM sodium chloride, 25 mM imidazole, 1 mM magnesium chloride, 0.1% (v/v) IGEPALCA-630 (Anatrace), 1 mM tris (2-carboxyethyl) phosphine (TCEP), 10% (v/v) glycerol with additional EDTA-free Protease Inhibitor Cocktail (Bimake), and then incubated with agitation for 20 min at 4 °C. The incubated cells were lysed with a high-pressure homogenizer operating at 500 bar. The supernatant was isolated by centrifugation at 30,000×g for 30 min, followed by incubation with Ni-NTA beads (GE Healthcare) for 2 h at 4 °C. After binding, the beads were washed with 20 column volumes of wash buffer of 25 mM HEPES, pH 7.4, 300 mM sodium chloride, 25 mM imidazole, 1 mM magnesium chloride, 1 mM TCEP, and 10% (v/v) glycerol. The protein was eluted with 3-4 column volumes of elution buffer of 25 mM HEPES pH 7.4, 300 mM sodium chloride, 300 mM imidazole, 1 mM magnesium chloride, 1 mM TCEP, and 10% (v/v) glycerol.

The co-expressed RdRp complex was incubated with additional nsp7 and nsp8 from the bacterial expression in a 1:1:2 molar ratios and incubated at 4 °C for 4h. Incubated RdRp complex was concentrated with a 100 kDa molecular weight cut-off centrifugal filter unit (Millipore Corporation) and then size-separated by a Superdex 200 Increase 10/300 GL column in 25 mM HEPES pH 7.4, 300 mM sodium chloride, 1 mM magnesium chloride, 1 mM TCEP. The fractions for the monomeric complex were collected and concentrated to up to 12 mg/ml. Suramin sodium salt (purchased from MedChemExpress) was dissolved in water up to 50 mM. Suramin derivatives (purchased from TopScience Co., Ltd) was dissolved in water at concentrations of 5 to 50 mM. For the suramin bound complex, the concentrated RdRp complex at the concentration of 12 mg/ml were incubated with 0.8 mM suramin at 4 °C for 0.5h for the next step of EM experiments.

#### Cryo-EM sample preparation and data acquisition

An aliquot of 3 μL protein sample of suramin-bound complex (12 mg/mL) containing 0.0035% DDM was applied onto a glow-discharged 200 mesh grid (Quantifoil R1.2/1.3), blotted with filter paper for 3.0 s and plunge-frozen in liquid ethane using a FEI Vitrobot Mark IV. Cryo-EM micrographs were collected on a 300kV Titan Krios microscope (FEI) equipped with a Gatan image filter (operated with a slit width of 20 eV) (GIF) and K3 direct detection camera. The microscope was operated at a calibrated magnification of 46,773 X, yielding a pixel size of 1.069 Å on micrographs. 11,846 micrographs in total were collected at an electron dose rate of 22.7 e^−^/Å^2^•s with a defocus range of −0.5 μm to −2.0 μm. Each movie with an accumulated dose of 68 e^−^/Å^2^ on sample were fractionated into a movie stack of 36 image frames.

#### Image processing

Frames in each movie stack were aligned for beam-induced motion correction using the program MotionCorr2(Zheng et al., 2017). CTFFIND4(Rohou and Grigorieff, 2015) was used to determine the contrast transfer function (CTF) parameters. 10,241 good micrographs were selected for further data processing. Auto-picking program of Relion 3.0(Zivanov et al., 2018) was used to pick the particles with the model of the apo RdRp complex of COVID-19 (PDB ID: 7BV1)(Yin et al., 2020) as a reference, yielding a total of 8,557,180 picked particles. Then, the extracted particle stack was transferred to software Cryosparc v2(Punjani et al., 2017) and a round of reference-free 2D classification was performed. 3,159,808 particles were selected from classes representing projections of suramin-bound RdRp complex in different orientations, and subjected to two rounds of Heterogenous Refinement using a reconstruction of the apo RdRp complex of COVID-19 (EMD-30209)(Yin et al., 2020) as a starting map. One converged 3D class with high-resolution feature contains one nsp12, one nsp7 and two copies of nsp8. The particles from that 3D class were then imported back into Relion 3.0 and subjected to a round of focused alignment with a mask including the whole protein components. Finally, 95,845 particles from a 3D class showing the highest resolution feature were selected for a round of 3D refinement. After a round of CTF refinement and Bayesian polishing of particles, another round 3D refinement was performed, yielding a final reconstruction at a global resolution of 2.57 Å based on the gold-standard Fourier shell correlation (FSC) = 0.143 criterion(Rosenthal and Henderson, 2003). The local resolution was calculated with Relion 3.0.

#### Model building

The model of suramin-bound RdRp complex was built by docking the model of apo structure of COVID-19 RdRp (PDB ID:7BV1) into the density map using UCSF Chimera(Pettersen et al., 2004), followed by *ab initio* model building of the N-terminal NiRAN domain of nsp12 and one copy of nsp8 in COOT(Emsley and Cowtan, 2004), and real space refinement using real_space_refine program in PHENIX(Adams et al., 2010). The model statistics were calculated with MolProbity(Chen et al., 2010) and listed in Supplemental Table S1. Structural figures were prepared in Chimera or ChimeraX(Goddard et al., 2018).

#### Preparation of template-primer RNA for polymerase assays

For the poly-A template-primer RNA, a short RNA oligonucleotide with sequence of 5’-FAM-GCUAUGUGAGAUUAAGUUAU-3’ (Sangon Biotech Co., Ltd) was used as the primer strand and a longer RNA oligonucleotide with sequence of 5’-AAAAAAAAAAAUAACUUAAUCUCACAUAGC -3’ (Sangon Biotech Co., Ltd) was used as template strand. To anneal the RNA duplex, both oligonucleotides were mixed at equal molar ratio in annealing buffer (10 mM Tris-HCl, pH 8.0, 25 mM NaCl and 2.5 mM EDTA), denatured by heating to 94 °C for 5 min and then slowly cooled to room temperature. The poly-U template- primer RNA was prepared similar to poly-A with the sequences of 5’-FAM-GCUAUGUGAGAUUAAGUUAU-3’ and 5’-UUUUUUUUUUAUAACUUAAUCUCACAUAGC -3’.

#### Gel mobility shift assay to detect RNA–RdRp protein binding

A gel mobility shift assay was performed to detect the effect of tested compounds on RNA binding by the RdRp complex. The binding reaction contained 20mM Tris-HCl 8.0, 10mM KCl, 6mM MgCl_2_, 0.01% Triton-X100, 1mM DTT, 1.14 U/ul RNase Inhibitor(Vazyme Biotech Co.,Ltd), 9 μg RdRp complex protein with1 μg poly-A template-primer RNA, and increasing amounts of corresponding compounds (0, 1, 10, 100, 1000 and 5000 μM for suramin, while 0, 1, 10, 100, 1000, 5000 and 10000 μM for RDV-TP). Binding reactions were incubated for 30 min at room temperature and resolved on 4-20% native polyacrylamide gel (Thermo Fisher) running in 1×TBE buffer at 90 V for 1.5 h in 4 °C cool-room. The gel was imaged with a Tanon-5200 Multi Fluorescence Imager according to the manufacturer’s protocol.

#### RdRp enzymatic activity assay and its inhibition by suramin

The purified SARS-CoV-2 RdRp complex from insect cell at final concentration of 1 μM was incubated with 3.0 μM poly-A template-primer RNA and 10 mM UTP(Macklin) in the presence of 1.14 U/μl RNase inhibitor in reaction buffer containing 20 mM Tris, pH8.0, 10 mM KCl, 6mM MgCl_2_, 0.01% Triton-X100, and 1mM DTT, which were prepared with DEPC-treated water. The total reaction volume was 20 μl. After incubation for 60 min at a 37°C water bath, 40μl quench buffer (94% formamide, 30 mM EDTA, prepare with DEPC-treated water) was added to stop the reaction. A sample of 18 μl of reaction was mixed with 2 μl of 10x DNA loading buffer (Solarbio). Half of the sample (10 μl) was loaded onto a 10% Urea-PAGE denatured gel, run at 120 V for 1h, and imaged with a Tanon-5200 Multi Fluorescence Imager. The setup for the inhibition assays of the RdRp by suramin is identical to the above for the RdRp enzymatic assays, except that suramin was added to final concentrations of 0, 1, 2, 4, 8, 16, 32 μM for 60 min before the addition of 10 mM UTP.

#### Fluorescence-based activity assay for SARS-CoV-2 RdRp

The detection of RNA synthesis by SARS-CoV-2 RdRp were established based on a real-time assay with the fluorescent dye SYTO 9 (Thermo Fisher), which binds dsRNA but not ssRNA template molecules. The fluorescence emitted was recorded in real-time using a TECAN F200 with excitation and emission filters at 485 and 520 nm, respectively. The assay records the synthesis of dsRNA in a reaction using a poly-U molecule as a template and ATP as the nucleotide substrate, which has been adapted from methods previously documented for the detection of Zika virus polymerase activity(Saez-Alvarez et al., 2019). Reactions were performed in individual wells of black 384-well low volume round bottom plates. The standard reaction contained 20 mM Tris-HCl, pH 8.0, 10 mM KCl, 6mM MgCl_2_, 180 μM ATP, 0.2μM poly-U template-primer RNA, 0.01%Trition-X100,1 mM DTT,0.025 U/ml RNase inhibitor (Vazyme Biotech Co., Ltd), and 0.25 μM SYTO 9 (50 μM stock solution in TE buffer pH 7.5). The assay was initiated by the addition of 5 μg/ml SARS-CoV-2 RdRp and the fluorescence was recorded over 30 min at room temperature. The reaction with equivalent of DMSO was set as MAX control, while the reaction with no SARS-CoV-2 RdRp was set as MIN control. The reactions were carried out in the presence of 0.2 μM poly-U template-primer RNA and 180 μM ATP, and increasing concentrations of each inhibitor. Fluorometric results were expressed as mean ± SD. Statistical significance was analyzed by two-way ANOVA using GraphPad Prism, version 8, as specified in the figure legends. Km determinations were obtained by plotting the velocity of the reaction as a function of nucleotide or ssRNA template concentrations using nonlinear regression. IC_50_ values were obtained by fitting the velocity data to a four-parameter logistic equation. Kinetic parameters and IC_50_ values were calculated using Sigmaplot, version 11.

#### Vero E6 cell based antiviral assay for suramin and suramin derivatives

African green monkey kidney Vero E6 cell line was obtained from American Type Culture Collection (ATCC, no. 1586) and maintained in Dulbecco’s Modified Eagle Medium (DMEM; Gibco Invitrogen) supplemented with 10% fetal bovine serum (FBS; Gibco Invitrogen), 1% antibiotic/antimycotic (Gibco Invitrogen), at 37 °C in a humidified 5% CO_2_ incubator. A clinical isolate of SARS-CoV-2 (nCoV-2019BetaCoV/Wuhan/WIV04/2019) was propagated in Vero E6 cells, and viral titer was determined as described previously(Wang et al., 2020). All the infection experiments were performed at biosafety level-3 (BSL-3).

Pre-seeded Vero E6 cells (5×10^4^ cells/well) were incubated with the different concentration of the indicated compounds for 1 hour, and then were infected with SARS-CoV-2 at a MOI of 0.01. Two hours later, the virus-drug mixture was removed and cells were further cultured with a fresh compound containing medium. At 24 h p.i.,., we measured viral RNA copy number in cell supernatant using real time PCR, as described previously(Wang et al., 2020). DMSO was used in the controls. At least two independent experiments were carried out for each compound.

#### The cytotoxicity assay

The cytotoxicity of the tested drugs on Vero E6 were determined by CCK8 assays (Beyotime, China).

#### Statistical analysis

The IC_50_ and EC_50_ values were expressed as mean ± SD from three independent experiments and determined via the nonlinear regression analysis using GraphPad Prism software 8.0 (GraphPad Software, Inc., San Diego, CA, USA)

### Supplemental Figures

**Figure S1.**
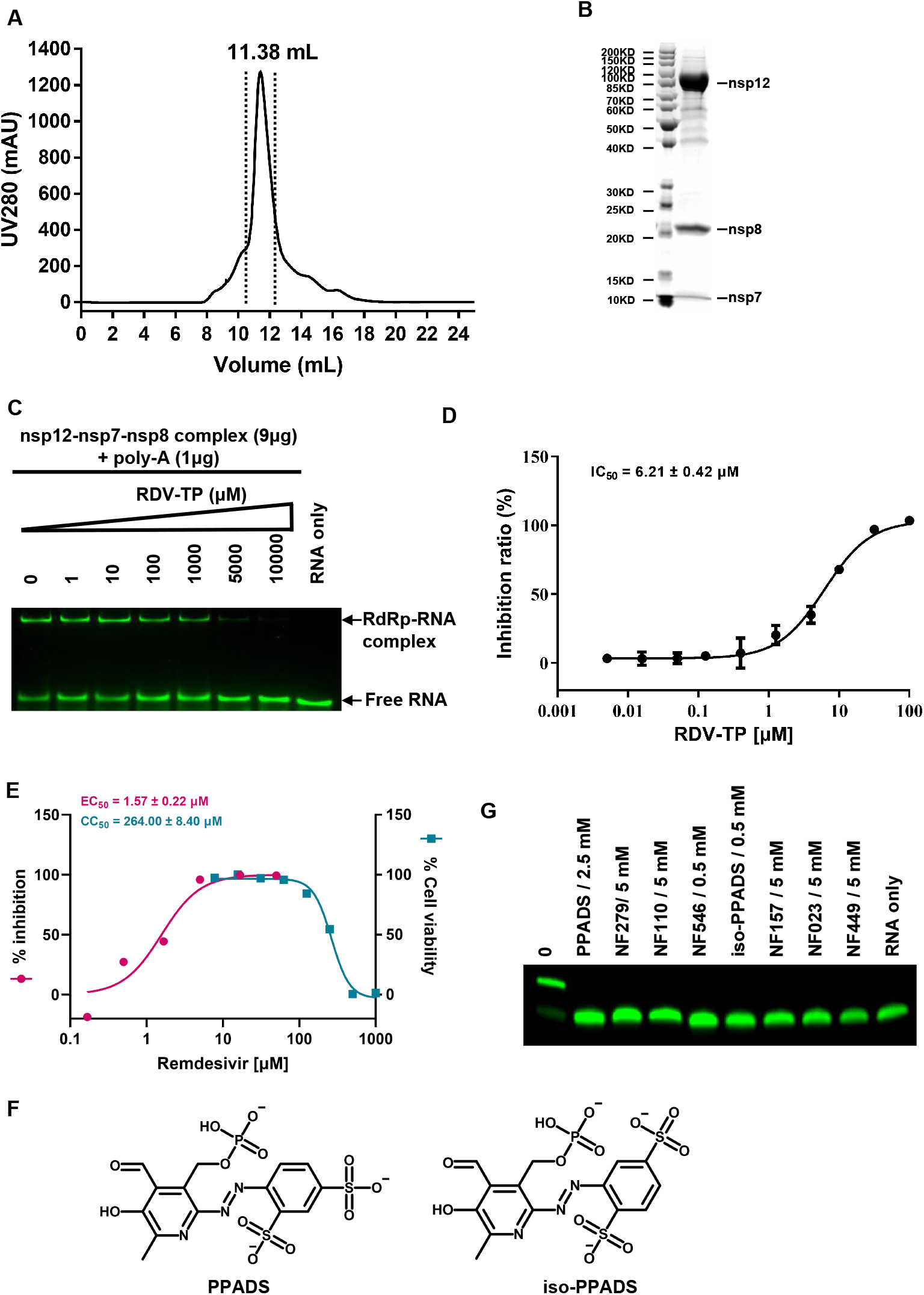
Purification and characterization of the RdRp complex. A. Gel filtration profile of the RdRp complex with additional nsp7 and nsp8, showing a sharp peak. B. SDS gel of the purified RdRp complex with additional nsp7 and nsp8, showing balanced ratios for each subunit. C. Gel mobility shift of the RdRp-RNA complex and the effect of RDV-TP. D. IC_50_ of RDV-TP for the RdRp complex. E. EC50 of remdesivir for SARS-CoV-2 inhibition and CC50 of remdesivir for cell-based toxicity. F. The structures of PPADS and iso-PPADS. G. Elongation of partial RNA duplex by the purified RdRp complex and its inhibition by 9 suramin derivatives at 0.5-5.0 mM concentrations depending on the solubility of each compound.

**Figure S2.**
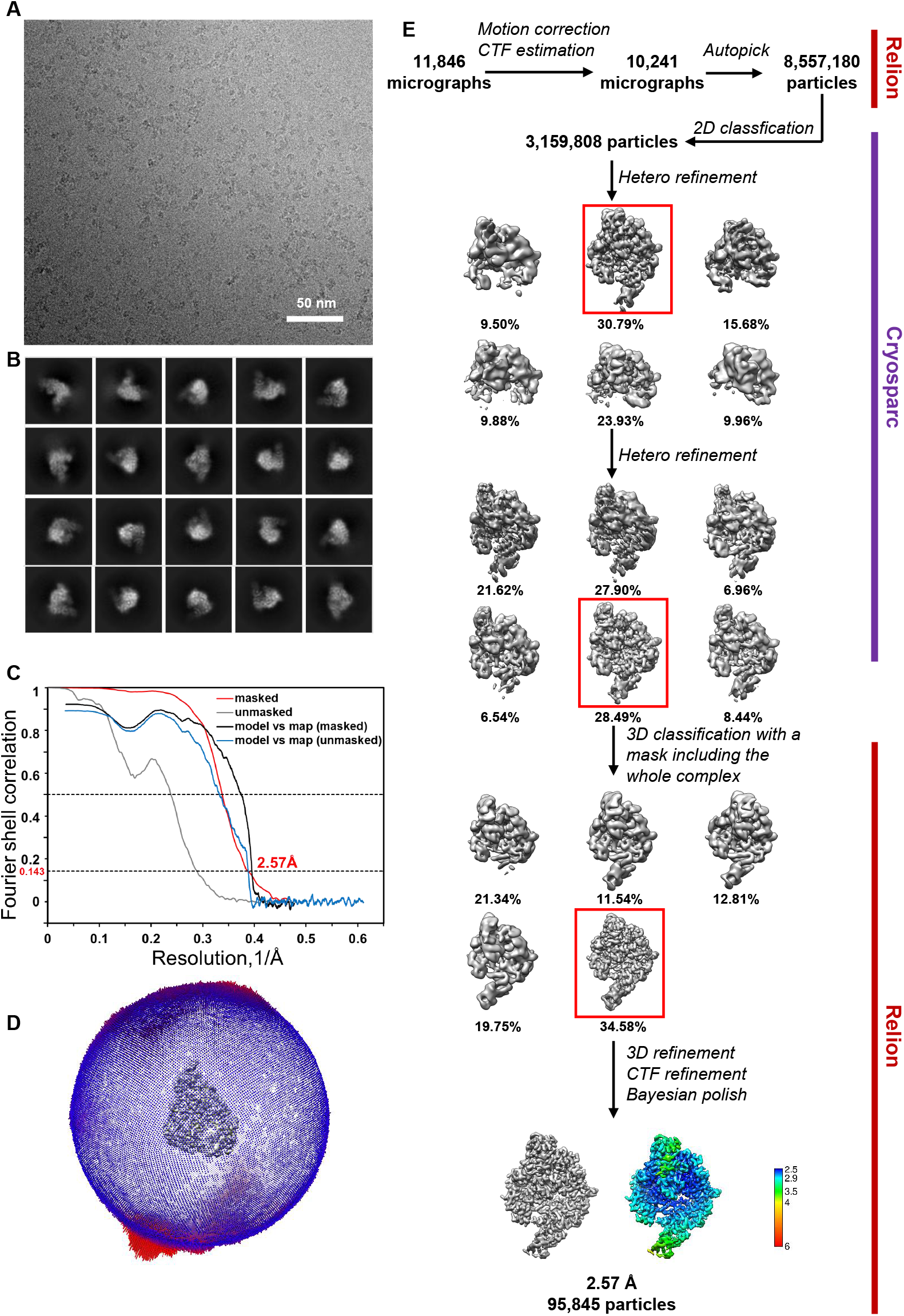
Single particle cryo-EM analysis of the RdRp-suramin complex. A. Representative cryo-EM micrograph of the RdRp-suramin complex. B. Representative 2D class averages of the RdRp-suramin complex. C. Fourier shell correlation curves of cryo-EM map for the the suramin-RdRp complex. D. Euler angle distribution of particles used in the final reconstruction. E. Flowchart of cryo-EM works of the suramin-RdRp complex with maps colored by local resolution (Å).

**Figure S3.**
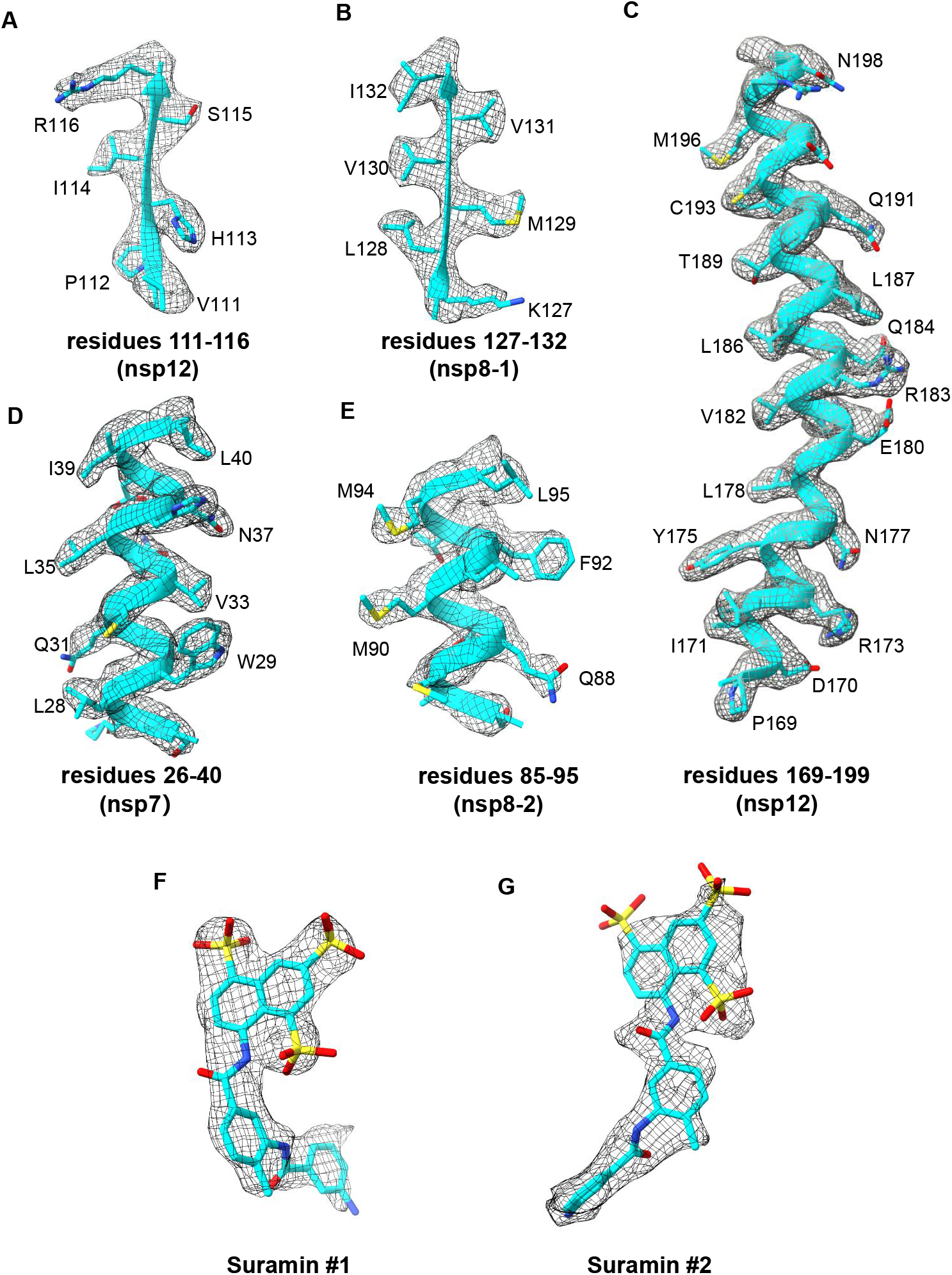
Cryo-EM map for the RdRp in complex with suramin. A-F. Cryo-EM map and model of nsp12 (residues 111-116) (A), nsp8-1 (residues 127-132) (B), nsp12 (residues 169-199) (C), nsp7 (residues 26-40) (D), nsp8-2 (residues 85–95) (E), and the two bound suramin molecules (F and G).

**Figure S4.**
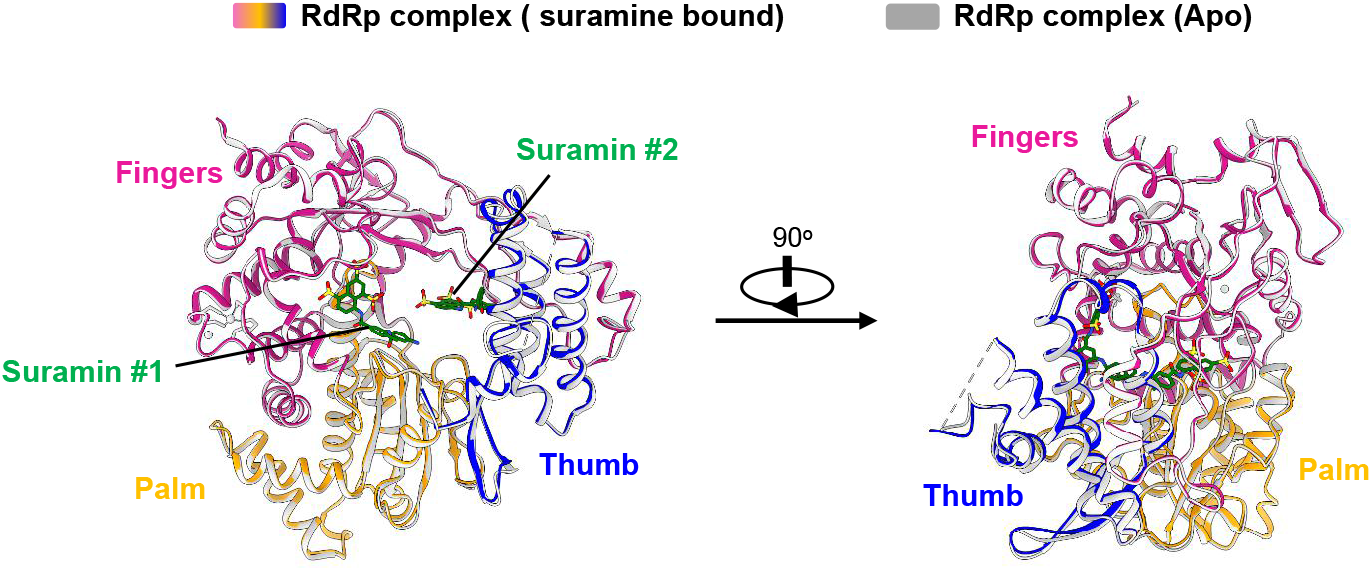
The structural comparison of the RdRp-suramin complex with the apo RdRp complex. The two overall views of the RdRp-suramin complex overlapped with the apo RdRp structure (PDB ID: 7BV1). For clarity, only the polymerase domains are retained. The apo RdRp structure is set as light gray.

**Figure S5.**
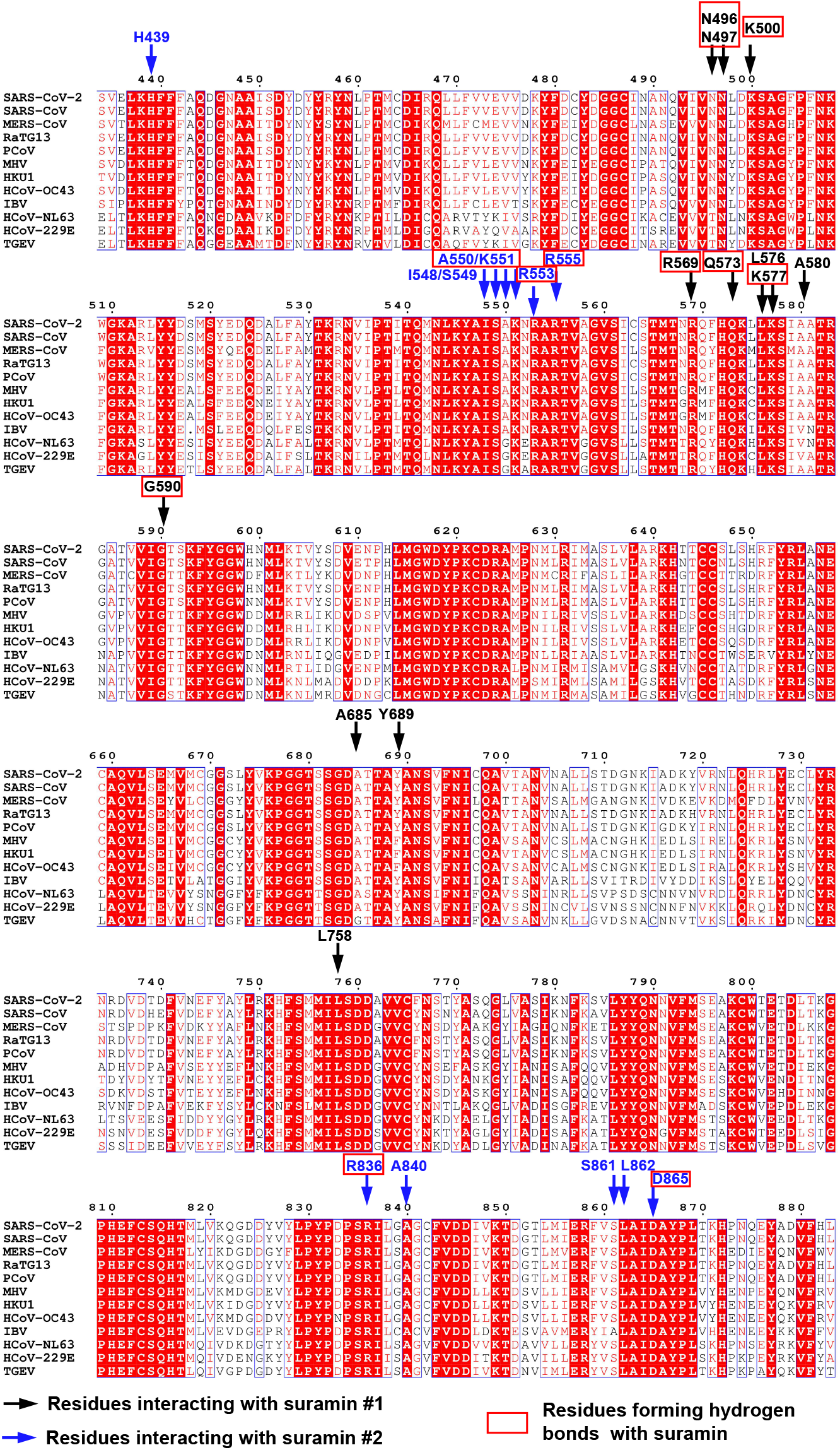
Sequence alignment of 12 coronavirus nsp12. Sequence alignment of nsp12 from eight β-CoVs (SARS-CoV-2, SARS-CoV, MERS-CoV, RaTG13, PCoV, MHV, HKU1, HCoV-OC43), one γ-CoVs (IBV) and three α-CoVs (HCoV-NL63, HCoV-229E, TGEV). Black arrows mark the residues interacting with suramin #1, blue arrows mark the residues interacting with suramin #2, while the red box sign the residues interacting with suramin molecules by hydrogen bonds.

**Figure S6.**
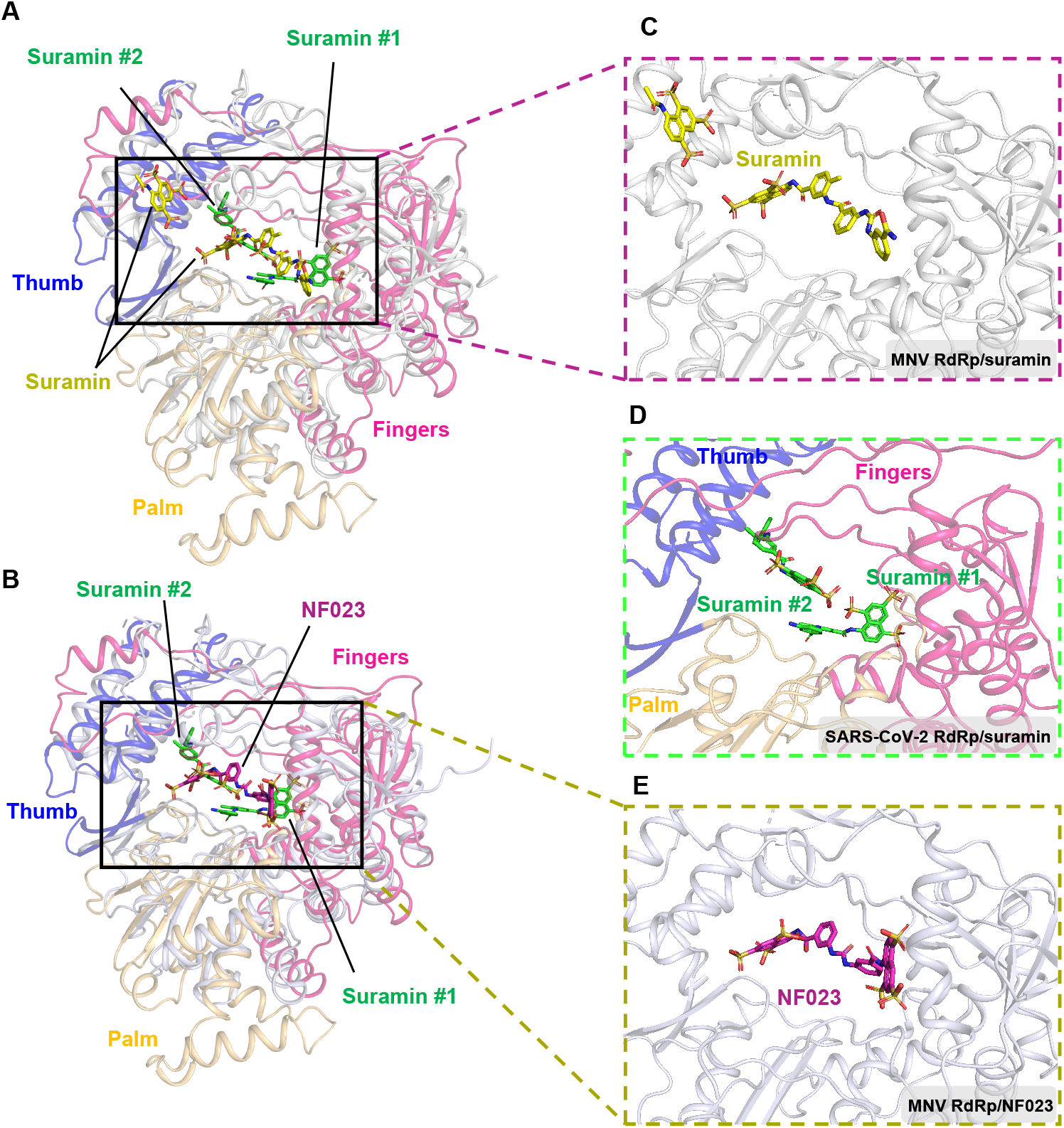
Comparison of SARS-CoV-2 RdRp-suramin complex with MNV (Murine Noroviruses) RdRp-suramin complex (PDB ID: 3UR0) and MNV RdRp-NF023(PDB ID: 3URF). A. Superimposition of the SARS-CoV-2 RdRp-suramin structure with the MNV RdRp-suramin structure based on the polymerase domain. Only the polymerase domain of SARS-CoV-2 RdRp is retained. B. Superimposition of SARS-CoV-2 RdRp-suramin structure with MNV RdRp-NF023 structure based on the polymerase domain. C. A close view of the suramin within the catalytic site in the MNV RdRp-suramin structure. D. A close view of the suramin within the catalytic site in the SARS-CoV-2 RdRp-suramin structure. E. A close view of the NF023 within the catalytic site in the MNV RdRp-NF023 structure.

**Figure S7.**
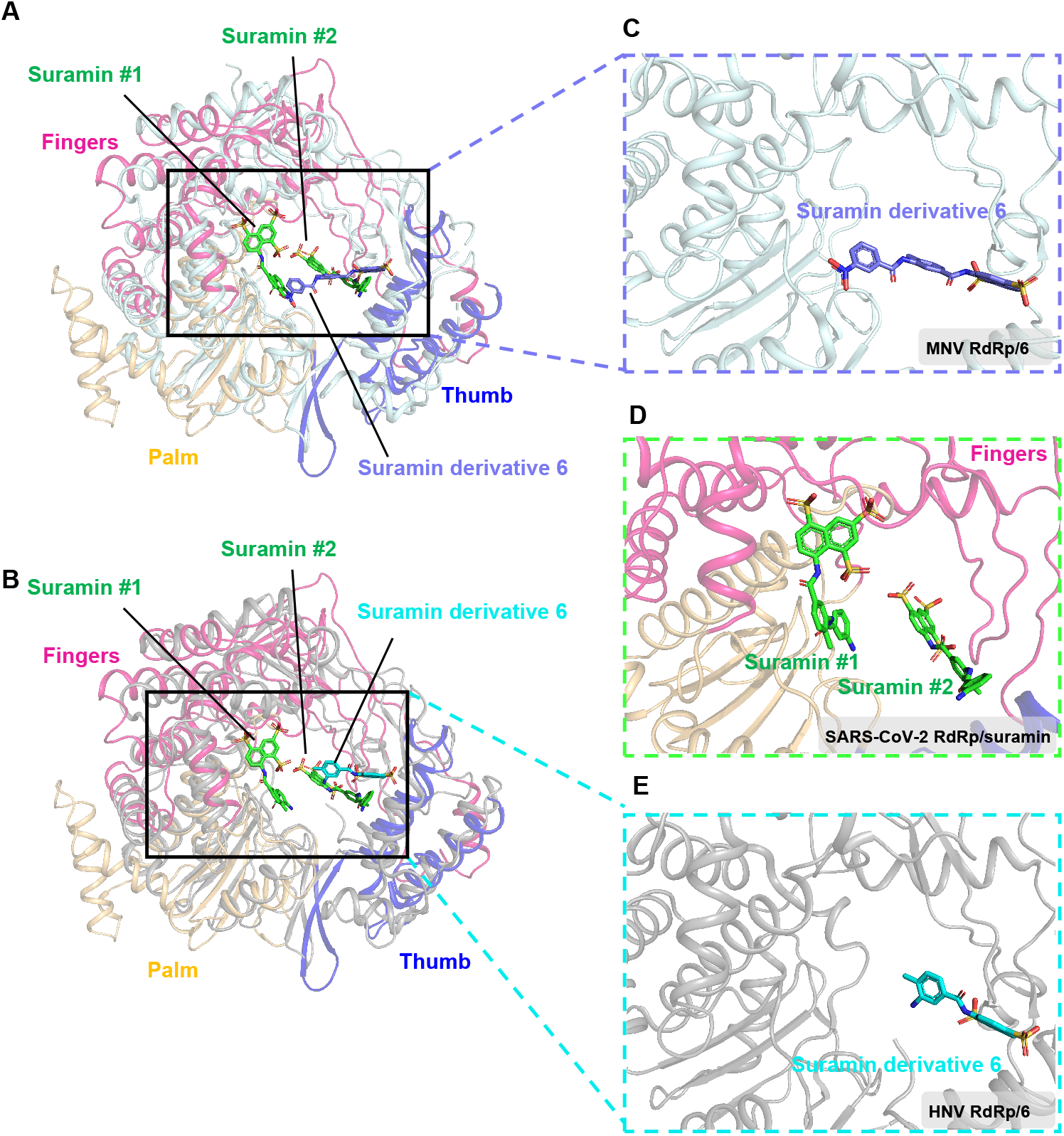
Comparison of SARS-CoV-2 RdRp-suramin complex with MNV RdRp-suramin derivative 6 complex (PDB ID: 4NUR) and HNV (Human Noroviruses) RdRp-suramin derivative 6 (PDB ID: 4NRT). A. Superimposition of SARS-CoV-2 RdRp-suramin structure with MNV RdRp-suramin derivative 6 structure based on the polymerase domain. Only the polymerase domain of SARS-CoV-2 RdRp is retained. B. Superimposition of SARS-CoV-2 RdRp-suramin structure with HNV RdRp-suramin derivative 6 structure based on the polymerase domain. C. A close view of the suramin derivative 6 within the active site in the MNV RdRp-suramin derivative 6 structure. D. A close view of the suramin within the active site in the SARS-CoV-2 RdRp-suramin structure. E. A close view of the suramin derivative 6 within the active site in the HNV RdRp-suramin derivative 6 structure.

**Supplemental Table S1.** Cryo-EM data collection, refinement, and validation statistics.

|  | suramin |
| --- | --- |
| **Data collection and Processing (for each dataset)** |  |
| Microscope | Titan Krios |
| Voltage (keV) | 300 |
| Camera | Gatan K3™ Camera |
| Magnification | 46,773 |
| Pixel size at detector (Å/pixel) | 1.069 |
| Total electron exposure (e-/Å<sup>2</sup>) | 68 |
| Exposure rate (e-/pixel/sec) | 26 |
| Number of frames collected during exposure | 36 |
| Defocus range (µm) | -0.5 to -2.0 |
| Phase plate (if used) | N/A |
| -phase shift range (in degrees) | N/A |
| -number of images per phase plate position | N/A |
| Automation software (EPU, SerialEM or manual) | SerialEM |
| Tilt angle (if grid was tilted) | 0 |
| Energy filter slit width (if used) | 20 |
| Micrographs collected (no.) | 11,846 |
| Micrographs used (no.) | 10,241 |
| Total extracted particles (no.) | 8,557,180 |
| **For each reconstruction:** |  |
| Refined particles (no.) | 95,845 |
| Final particles (no.) | 95,845 |
| Point-group or helical symmetry parameters | C1 |
| Resolution (global, Å) |  |
| FSC 0.5 (unmasked/masked) | 4.2/3.5 |
| FSC 0.143 (unmasked/masked) | 3.0/2.6 |
| Resolution range (local, Å) | 2.5-6.0 |
| Map sharpening B factor (Å<sup>2</sup>) | -27.866 |
| **Model composition (for each model)** |  |
| Protein | 1181 |
| Ligands | 4 |
| RNA/DNA | 0 |
| **Model Refinement (for each model)** |  |
| Refinement package | PHENIX-1.14_3260 |
| -real or reciprocal space | real space |
| Initial model used (PDB code) | 7BV1 |
| Model-Map CC | 0.85 |
| Model resolution (Å) | 2.7 |
| FSC threshold | 0.5 |
| B factors (Å<sup>2</sup>) |  |
| Protein residues | 63.49 |
| Ligands | 64.64 |
| RNA/DNA | N/A |
| R.m.s. deviations from ideal values |  |
| Bond lengths (Å) | 0.012 |
| Bond angles (°) | 0.904 |
| **Validation (for each model)** |  |
| MolProbity score | 1.34 |
| CaBLAM outliers | 0.96 |
| Clashscore | 5.43 |
| Poor rotamers (%) | 0.76 |
| C-beta deviations | 0.00 |
| EMRinger score (if better than 4 Å resolution) | 3.63 |
| Ramachandran plot |  |
| Favored (%) | 97.77 |
| Outliers (%) | 0.00 |

